# Identification and characterization of the functional LolB ortholog in *Bacteroides*

**DOI:** 10.64898/2026.02.05.704107

**Authors:** Krista M. Armbruster, Rhea Trickannad, Nichollas E. Scott, Nicholas A. Pudlo, Jesse W. Wotring, Eric C. Martens, Jonathan Z. Sexton, Nicole M. Koropatkin

## Abstract

Bacteroidota are prolific members of the human gut microbiota, influencing overall health through the degradation of various polysaccharides. To aid in this process, these bacteria deploy cell surface lipoproteins, proteins anchored to the membrane by a lipidated N-terminal cysteine. Despite their importance, lipoprotein synthesis and transport in Bacteroidota is not well defined, particularly how lipoproteins reach the outer membrane and cell surface. In *Escherichia coli*, lipoproteins are inserted into the outer membrane by LolB, the final step of the Lol pathway for localizing lipoproteins, previously thought to be confined to γ- and β-proteobacteria. Herein, we report a structural ortholog of LolB in *Bacteroides* that rescues the lethal phenotype associated with LolB depletion in *E. coli*. We demonstrate this ortholog’s LolB-like activity through mutagenesis studies and a lipoprotein insertion activity assay. We find that the gene cannot be deleted from *Bacteroides*, but that depletion does not significantly impact growth on polysaccharides, nor the surface localization of lipoproteins involved in starch degradation. Nonetheless, depletion significantly alters the composition of many proteins in both the inner and outer membranes, while others remain unchanged. Altogether, our findings contribute to elucidating the lipoprotein transport pathway in *Bacteroides* and how it impacts cell physiology.

**Importance:** Bacteroidota are abundant members of the human gut microbiota that influence health and disease. These bacteria deploy numerous cell surface lipoproteins that mediate their interactions with the host and play key roles in cell physiology. However, their mechanism(s) of lipoprotein transport is understudied. Here, we used a genetic screen to identify orthologs to *E. coli* LolB, the protein responsible for lipoprotein insertion into the outer membrane. Our screen revealed a structural ortholog to LolB in *Bacteroides* that performs this function. We show that when LolB is depleted, lipoproteins still reach the cell surface, even though the overall protein composition of the membrane is significantly altered. Our results broaden the understanding of both lipoprotein transport and Bacteroidota physiology.

## Introduction

As prevalent members of the human gut microbiota, Bacteroidota heavily influence health and disease, largely through the degradation of diverse polysaccharides (1–3). Breakdown of glycans feeds neighboring gut inhabitants and provides beneficial byproducts to the host (4). This process typically requires one or more cell surface lipoproteins, a class of membrane proteins characterized by a lipidated N-terminal cysteine (5, 6). Beyond polysaccharide degradation, lipoproteins function in many other processes, including cell envelope integrity, antibiotic resistance, immunomodulation, virulence, and more (6–9).

Lipoproteins are initially targeted to the inner membrane (IM) as a precursor, featuring a signal peptide II (SPII) and a lipobox motif marking the protein for lipidation (10, 11). The precursor is sequentially modified by three enzymes (Lgt, Lsp, and Lnt) to ultimately lipidate the cysteine (12–14). The last enzyme in this pathway, lipoprotein *<u>N</u>*-acyltransferase (Lnt), attaches a lipid to the α-amino group of the cysteine to form the mature lipoprotein (14). Once *N*-acylated, lipoproteins enter the localization <u>o</u>f lipoprotein (Lol) export machinery, where those destined for the outer membrane (OM) are extracted from the IM by the ABC transporter LolCDE (15). The lipoprotein is transferred to LolA, a chaperone that shields the lipoprotein’s hydrophobic N-terminus as the LolA:lipoprotein complex transverses the periplasmic space (16). The complex docks with LolB, itself a lipoprotein, that is anchored to the inner leaflet of the OM (17). The lipoprotein is transferred from LolA to LolB in a mouth-to-mouth fashion (18), then inserted into the OM by an unknown mechanism.

Most knowledge of lipoprotein biosynthesis and transport has resulted from studies on the model diderm *Escherichia coli*. Empirical study of these processes in Bacteroidota was lacking until recently, which in turn revealed several differences from *E. coli*. In 2024, we showed that lipoprotein *N*-acylation is executed not by Lnt, but by a previously uncharacterized protein named lipoprotein *<u>N</u>*-acyltransferase in *<u>B</u>acteroides* (Lnb)(19). Lnb is distinct in sequence and structure from Lnt and is not essential for *Bacteroides* growth (19), whereas Lnt is essential in *E. coli* (20). Furthermore, Bacteroidota were thought to not encode LolB, since sequence orthologs of LolB are absent from their genomes (8, 21, 22). However, De Smet *et al*. identified candidate LolB-like proteins throughout Bacteroidota by *in silico* distant structural homolog prediction to *E. coli* LolB (23). Similar to Lnt/Lnb, LolB is essential in *E. coli* (24), but was found dispensable in *Flavobacterium johnsoniae* (23). Lastly, Bacteroidota localize many lipoproteins to the cell surface, but exactly how this occurs is unknown (22, 25). The current understanding of lipoprotein surface exposure in Bacteroidota is limited to the lipoprotein <u>e</u>xport <u>s</u>ignal (LES), a negatively charged consensus sequence immediately following the acylated cysteine of cell surface lipoproteins, that appears to dictate their localization (26, 27). It is even unclear if all lipoproteins move through the Lol system, if surface-destined lipoproteins move through a separate system, or a combination of the two.

To continue elucidating how lipoprotein transport occurs in Bacteroidota, we identified the functional LolB equivalent in *Bacteroides* by a complementation rescue assay in *E. coli*. We validate its LolB-like activity through a lipoprotein insertion assay and find that we cannot delete it from the native organism. Depletion of *Bacteroides* LolB minimally impacts growth on polysaccharides and the cell surface localization of lipoproteins involved in starch degradation. Depletion does, however, significantly alter the protein composition of both IM and OM while other proteins remain unchanged, supporting the existence of an alternate pathway for lipoprotein localization. Altogether, our findings broaden the repertoire of proteins involved in lipoprotein transport and explore the role of LolB in *Bacteroides* cell physiology.

## Results

### Discovery of a *Bacteroides* gene that complements LolB-depleted *E. coli*

Loss of LolB is fatal to *E. coli* due to deficiencies in lipoprotein trafficking (24). We reasoned this could be leveraged to identify functional orthologs of LolB from *Bacteroides* capable of restoring lipoprotein transport in LolB-depleted *E. coli*, as in previous similar studies (19, 28). To construct a conditionally lethal *lolB* mutant of *E. coli*, we integrated the Kan^R^-TT-*araC*-*P*_*BAD*_ cassette directly upstream of *lolB* (Fig. 1A). This was accomplished in an *lpp*-null background, where Lpp is a highly abundant OM lipoprotein in *E. coli* (29). The resulting strain KA38 largely depends on arabinose-induced expression of *lolB* via the *P*_*BAD*_ promoter for growth (Fig. 1B). Minimal growth was observed on glucose, which represses the *P*_*BAD*_ promoter and thus expression of *lolB*.

**Figure 1.**
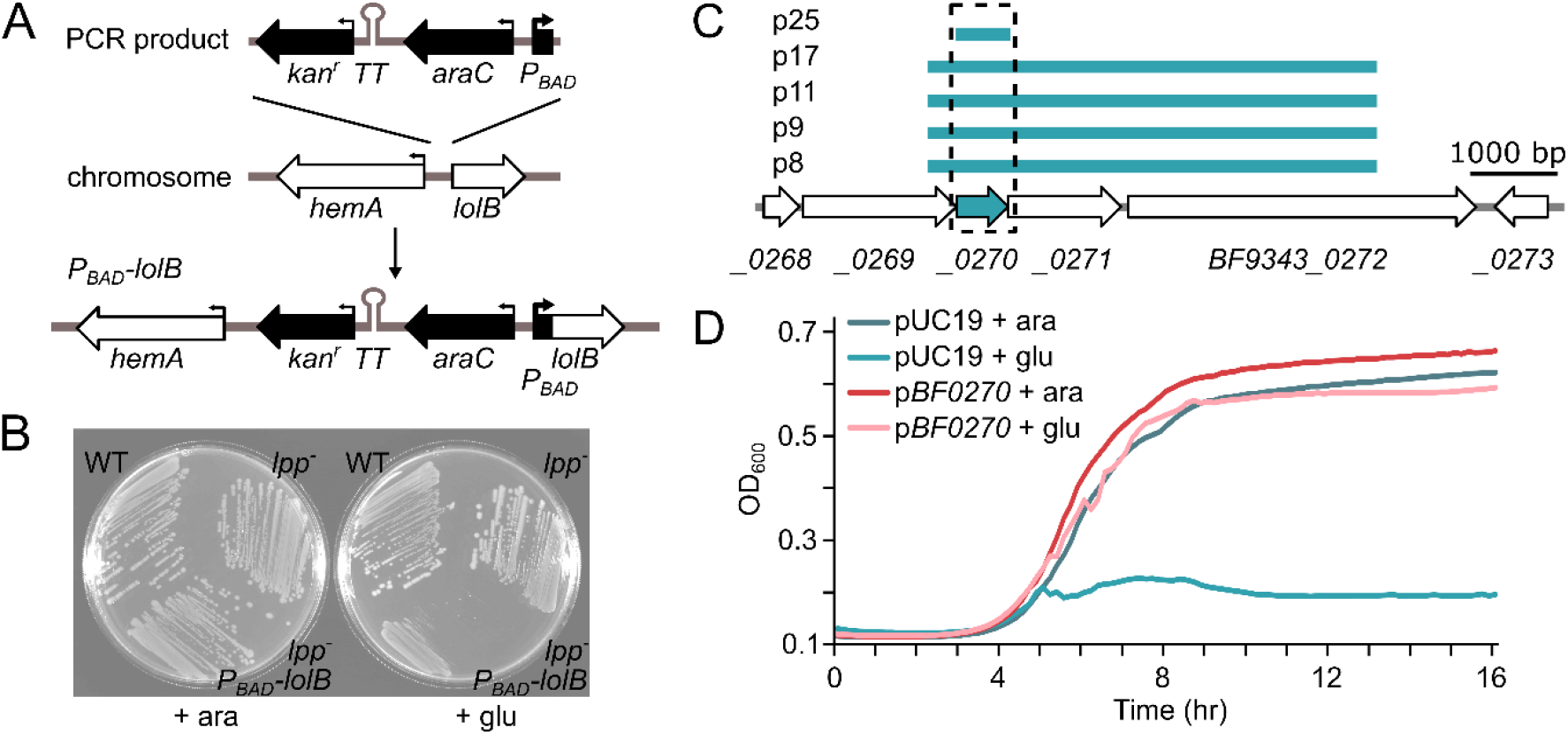
Identification of BF9343_0270 as the candidate *lolB* gene in *Bacteroides*. (A) The Kan^R^-TT-*araC*-*P*_*BAD*_ cassette was integrated directly upstream of *lolB* to generate the conditional lethal strain KA38. (B) Wildtype *E. coli* (WT), *lpp*^*-*^ (strain TXM327), and *lpp*^*-*^ *P*_*BAD*_*-lolB* (strain KA38) were struck onto LB agar plates containing arabinose (ara) or glucose (glu). (C) DNA fragments rescuing the growth of LolB-depleted KA38 cells were mapped to the *B. fragilis* genome, showing overlap at open reading frame *BF9343_0270* (boxed and shaded). (D) The growth of strain KA38 harboring either empty pUC19 or pUC19-*BF9343_0270* (shortened to p*BF0270*) with arabinose or glucose as measured by OD_600_ over time.

To screen for *Bacteroides* genes capable of rescuing LolB depletion, KA38 was transformed with a library of *Bacteroides fragilis* genomic DNA and plated onto solid media containing glucose. The same library identified Lnb in a similar experiment (19). Colonies that grew on glucose indicate phenotypic rescue, and multiple fragments of *B. fragilis* genomic DNA were recovered (Fig. 1C). The fragments overlap on a single open reading frame BF9343_0270, a predicted lipoprotein encoding the domain of unknown function (DUF) 4292, the same protein indicated as a putative LolB by De Smet *et al* (23). To confirm rescue, we cloned BF9343_0270 by itself and transformed it into KA38, then measured the growth of the resulting strain with arabinose and glucose compared to KA38 with pUC19 empty vector (Fig. 1D). The addition of BF9343_0270 permitted growth even with glucose, while KA38 with pUC19 could not grow without arabinose.

### The *Bacteroides* LolB candidate is structurally similar to *Ec*LolB

We validated our LolB-like candidate in *Bacteroides thetaiotaomicron* (*B. theta*), a close relative of *B. fragilis* that is genetically tractable and a model organism for studies in our lab. The *B. theta* homolog *BT_3463* is 59% identical (77% similar) to *BF9343_0270* and shares the same genomic neighborhood. We refer to this gene as *Bt*LolB going forward.

Despite sharing no sequence homology, a comparison of the crystal structure of *Ec*LolB (PDB 1IWN)(30) to the AlphaFold3-predicted (31) structure of *Bt*LolB reveals similarities (Fig. 2A). Both feature a concave antiparallel β-sheet (11 β-strands in *Ec*LolB; 8 β-strands predicted in *Bt*LolB) forming an open β-barrel that is covered by a three α-helical lid. This fold, described as a catcher’s mitt, is smaller in the predicted structure of *Bt*LolB (residues 40-197) than *Ec*LolB (residues 15-186). *Bt*LolB is also predicted to have an unstructured N-terminus (residues 27-39) after the SPII signal peptide instead of the N-terminal α-helix (residues 15-28) that shapes one side of the *Ec*LolB barrel. The interior of both proteins is highly hydrophobic, the apparent lipid-binding site of traversing lipoproteins.

**Figure 2.**
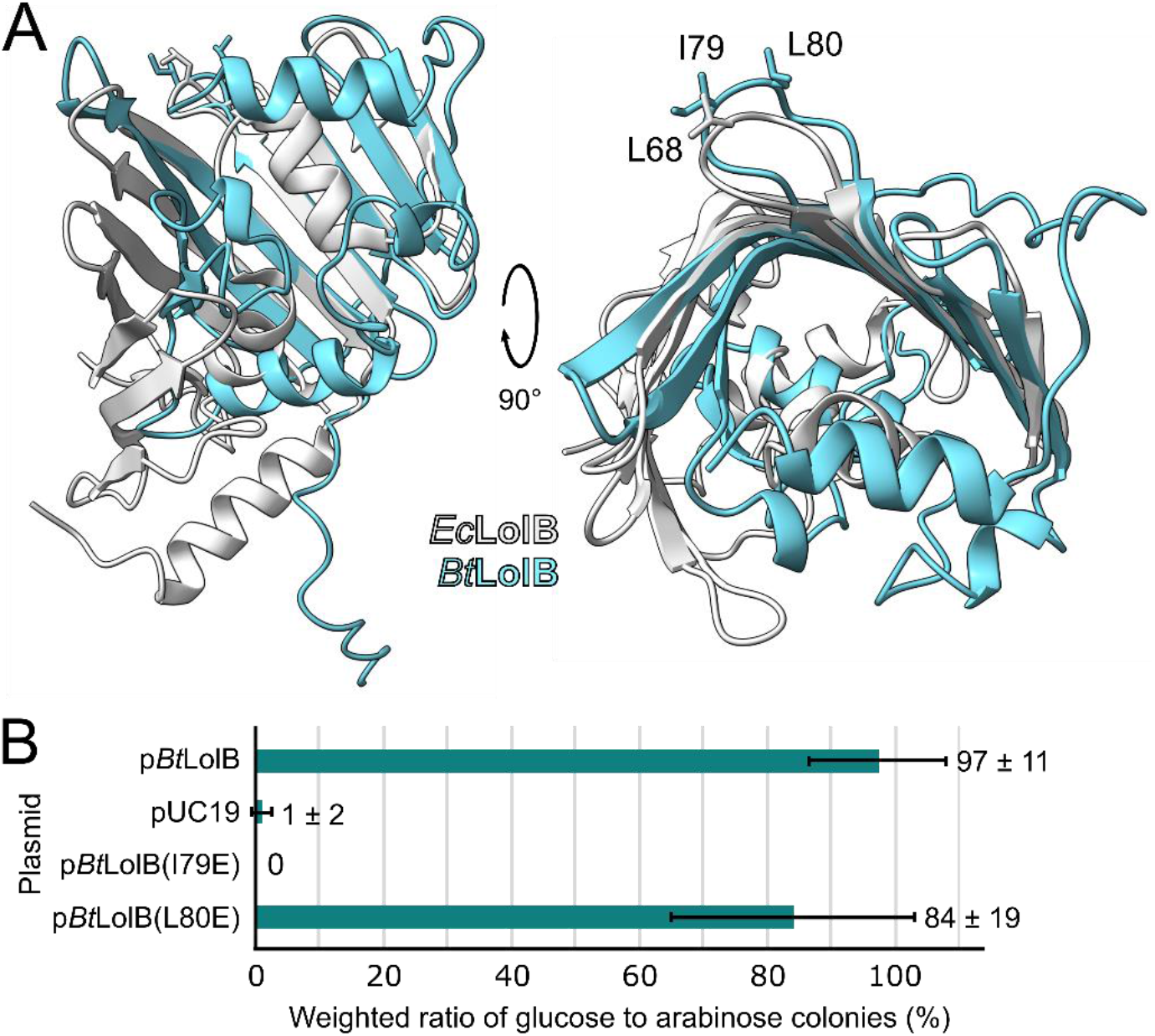
Structural comparisons and essential loop residue analysis of *Bt*LolB. (A) An overlay of the AlphaFold3-predicted structure of *Bt*LolB (aqua) to the solved structure of *Ec*LolB (white; PDB 1IWN). The residues L68 from *Ec*LolB and I79 and L80 from *Bt*LolB are indicated on the loop of interest. (B) KA38 (*lpp*^*-*^ *P*_*BAD*_*-lolB*) was transformed with the indicated plasmids and spread onto solid media containing arabinose or glucose. The resulting colonies are represented as a weighted ratio of glucose-grown to arabinose-grown colonies.

Previous work revealed the essential role of a protruding loop of *Ec*LolB in inserting lipoproteins into the OM (32). Mutation of L68 of this loop to acidic, but not hydrophobic, residues abrogates *Ec*LolB insertase activity. Based on the structural overlay of *Ec*LolB and *Bt*LolB, *Bt*LolB has a similar loop, with I79 and L80 overlapping with L68 of *Ec*LolB (Fig. 2A). To determine if either of these residues are essential to *Bt*LolB activity, we mutated each to glutamic acid and screened for rescue in KA38. Cells expressing *Bt*LolB(L80E) grew under *Ec*LolB-depleting conditions (Fig. 2B), though colonies were notably smaller than those expressing wildtype *Bt*LolB, suggesting reduced ability to complement *Ec*LolB depletion. Cells expressing *Bt*LolB(I79E) strikingly did not grow under *Ec*LolB-depleting conditions. These data suggest that *Bt*LolB functions similarly to *Ec*LolB, with I79 performing a crucial role in *Bt*LolB activity.

### *Bt*LolB exhibits membrane insertase activity

To determine if *Bt*LolB is a bona fide LolB-like insertase, we performed an *in vitro* assay measuring lipoprotein insertion into membranes, as done previously (32–35). First, we harvested OMs from wildtype *B. theta* and KA308, a strain with *Bt*LolB-Myc under the control of an anhydrotetracycline (aTc)-inducible promoter, grown with or without aTc. This strain was constructed as *Bt*LolB could not be deleted from the *B. theta* genome (described below). Next, we purified His-tagged *Ec*LolA from the periplasm of *E. coli* constitutively expressing Strep-tagged Lpp. LolA and Lpp form a soluble complex that is readily purified as a substrate for LolB (36). The LolA:Lpp complex was then incubated with the *B. theta* OMs. If *Bt*LolB has insertase activity, a band corresponding to Lpp-Strep would be observed by immunoblot in the OMs recovered from the reaction. Indeed, Lpp-Strep signal was detected in all samples where *Bt*LolB is present but was strikingly missing from the *Bt*LolB-depleted sample (Fig. 3A). A control blot against *Ec*LolA-His after OM recovery shows no contaminating LolA:Lpp complex that could account for Lpp-Strep signal (Fig. 3A). A Ponceau-stained membrane serves as a loading control (Supp. Fig. 1). These data also confirm our *in vivo* screen (Fig. 1), that *Ec*LolA works with *Bt*LolB.

**Figure 3.**
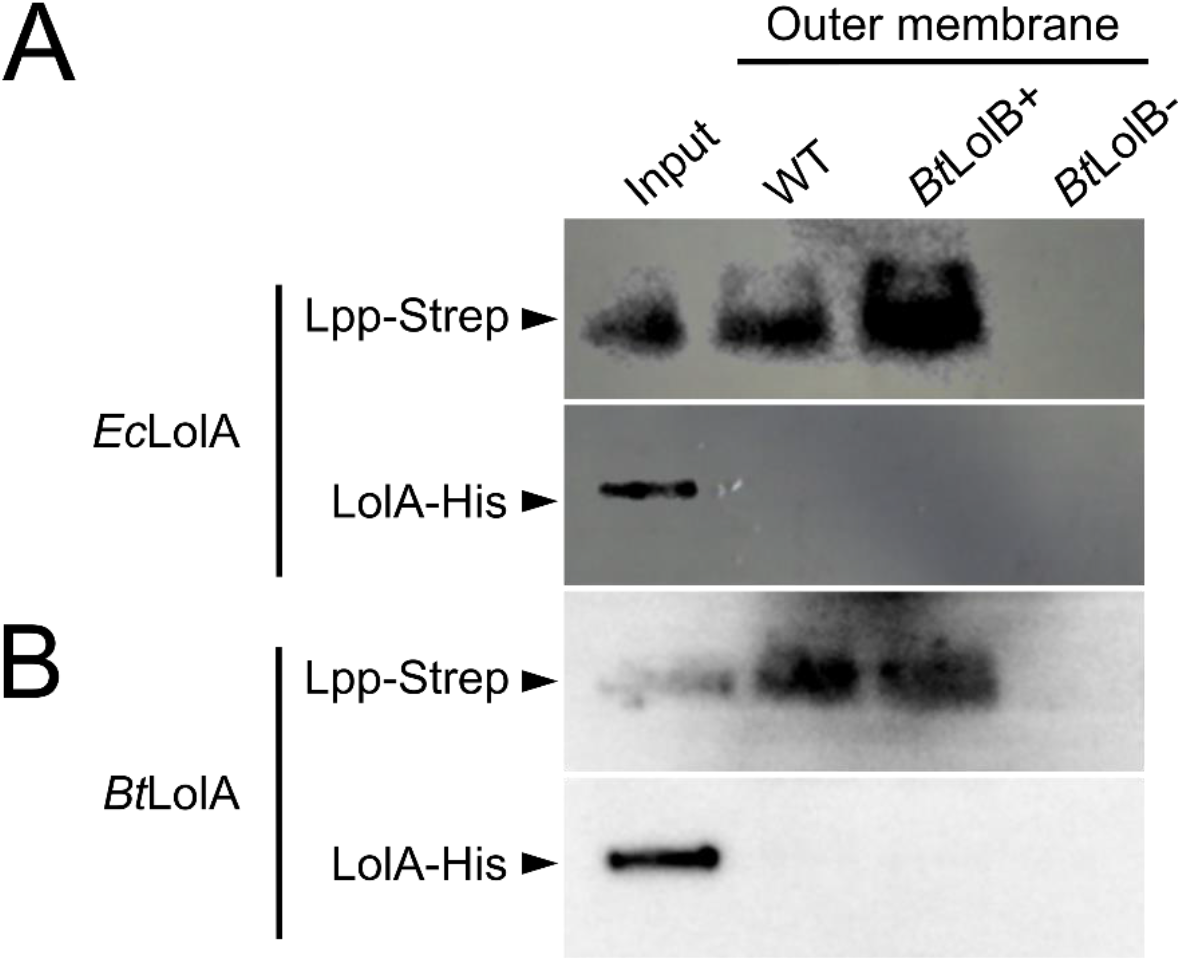
Evidence of LolB-like insertase activity by *Bt*LolB. (A) Immunoblots against Lpp-Strep and *Ec*LolA-His in OMs recovered from the reaction with *Ec*LolA:Lpp. (B) Immunoblots against Lpp-Strep and *Ec*LolA-His in OMs recovered from the reaction with *Bt*LolA:Lpp. For both, the “input” lane is the purified LolA:Lpp complex. *Bt*LolB+ and *Bt*LolB- refer to OMs of the *Bt*LolB-inducible strain KA308 grown with or without aTc, respectively.

Recent studies have described bifunctional LolA proteins with both chaperone and insertase activity (34, 37). To address this possibility in *B. theta*, we repeated the above experiment using the putative LolA from *B. theta* (BT_4335; inferred from homology to *Ec*LolA). Again, Lpp-Strep signal was detected only in samples where *Bt*LolB is present, suggesting that *Bt*LolA does not insert lipoproteins into the OM (Fig. 3B). Taken together, we conclude that *Bt*LolB is the true LolB of *B. theta*.

### *Bt*LolB associates with lipoproteins

To characterize lipoprotein association with *Bt*LolB within *B. theta*, we performed a co-immunoprecipitation (co-IP). Cells overexpressing *Bt*LolB-Myc from an *att* site (with the endogenous *Bt*LolB deleted; strain KA282) were cross-linked with formaldehyde. This seemed a necessary step to capture *Bt*LolB with its lipoprotein cargo, as the interaction between *Bt*LolB and lipoprotein is likely very transient (17, 30). *Bt*LolB-Myc was purified from OMs of cross-linked cells using anti-Myc beads and analyzed by liquid-chromatrography mass spectrometry. Paired samples were pulled down with anti-HA beads as isotype controls.

Using a log_2_ fold-change of ≥ 2.0 and *P* < 0.05 as determined by Student’s t-tests, our experiment identified eight proteins that co-purified with *Bt*LolB-Myc (Table 1). SignalP 6.0 predicts three of these proteins to have an SPII signal peptide, likely lipoproteins (38). However, upon further investigation, we predict six of the eight to be lipoproteins, based on previous publications or the following criteria: i) a cysteine located 15-35 amino acids into the protein sequence, ii) the three amino acids preceding the cysteine are largely uncharged and nonpolar, to echo the lipobox motifs defined for phyla other than Bacteroidota (10, 11), and iii) a disordered N-terminal region common to lipoproteins (39), as predicted by AlphaFoldDB (40). Our interpretation of this data supports *Bt*LolB association with lipoproteins in the OM of *B. theta*. The full co-IP results are provided in Supp. Dataset 1.

**Table 1.**
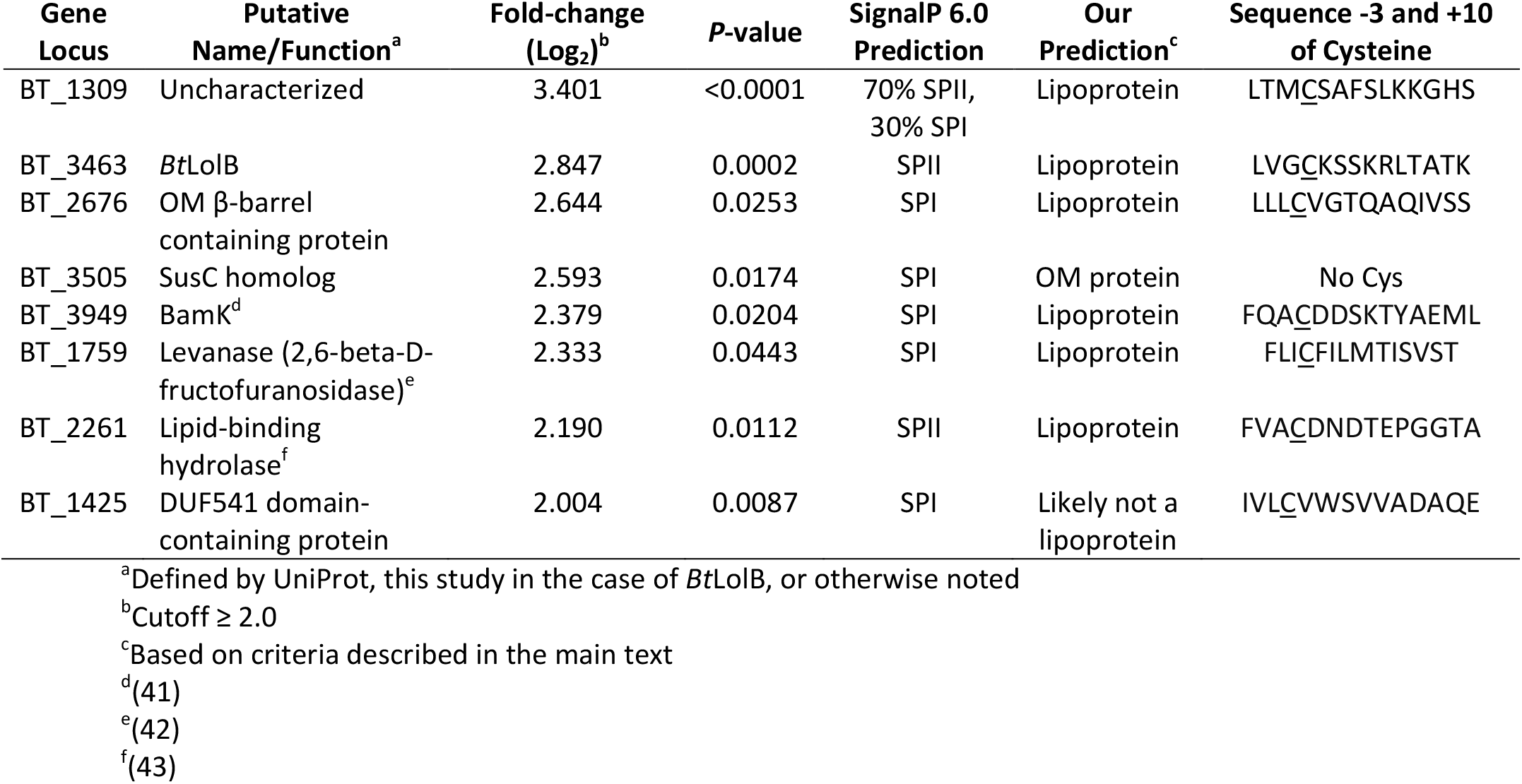
*Bt*LolB-Myc co-immunoprecipitation enriched proteins.

| Gene Locus | Putative Name/Function<sup>a</sup> | Fold-change (Log<sub>2</sub>)<sup>b</sup> | *P*-value | SignalP 6.0 Prediction | Our Prediction<sup>c</sup> | Sequence -3 and +10 of Cysteine |
| --- | --- | --- | --- | --- | --- | --- |
| BT_1309 | Uncharacterized | 3.401 | <0.0001 | 70% SPII, 30% SPI | Lipoprotein | LTMCSAFSLKKGHS |
| BT_3463 | *BtLolB* | 2.847 | 0.0002 | SPII | Lipoprotein | LVGCKSSKRLTATK |
| BT_2676 | OM β-barrel containing protein | 2.644 | 0.0253 | SPI | Lipoprotein | LLLCVGTQAQIVSS |
| BT_3505 | SusC homolog | 2.593 | 0.0174 | SPI | OM protein | No Cys |
| BT_3949 | BamK<sup>d</sup> | 2.379 | 0.0204 | SPI | Lipoprotein | FQACDDSKTYAEML |
| BT_1759 | Levanase (2,6-beta-D-fructofuranosidase)<sup>e</sup> | 2.333 | 0.0443 | SPI | Lipoprotein | FLICFILMTISVST |
| BT_2261 | Lipid-binding hydrolase<sup>f</sup> | 2.190 | 0.0112 | SPII | Lipoprotein | FVACDNDTEPGGTA |
| BT_1425 | DUF541 domain-containing protein | 2.004 | 0.0087 | SPI | Likely not a lipoprotein | IVLCVWSVVADAQE |

<sup>a</sup>Defined by UniProt, this study in the case of *BtLolB*, or otherwise noted
<sup>b</sup>Cutoff ≥ 2.0
<sup>c</sup>Based on criteria described in the main text
<sup>d</sup>(41)
<sup>e</sup>(42)
<sup>f</sup>(43)

### *Bt*LolB-depleted cells grow on various carbon sources

Multiple transposon studies have assigned *Bt*LolB as essential (44, 45). To instead study the effects of *Bt*LolB depletion, we constructed strain KA308 where *BtlolB-myc* was integrated into an *att* site under the control of an aTc-inducible promoter (46), then the native *BtlolB* was deleted from the genome. We hypothesized that this strain would not grow without inducer; however, experiments revealed robust growth even without aTc in both rich media (tryptone-yeast extract-glucose; TYG) and minimal media (MM) with glucose (Supp. Fig. 2A, Fig. 4A). Analysis by RT-qPCR showed a significant average of 18.9-fold reduction of *BtlolB-myc* expression without aTc compared to with aTc, and transcript levels of *BtlolB-myc* with aTc are comparable to that of wildtype (Supp. Fig. 3).

*Bacteroides* spp. are known for their ability to uptake and degrade polysaccharides as carbon sources using cell surface lipoproteins (2). In our previous study, deletion of Lnb resulted in growth defects that varied depending on the carbon source (19), and we wondered if the same would be true upon depletion of *Bt*LolB. To test this, we grew our *Bt*LolB-inducible strain in MM supplemented with a panel of carbohydrates, with or without aTc inducer. We also included Δ*lnb* cells for a direct comparison. Addition of aTc to wildtype or Δ*lnb* cells did not significantly impact growth (Supp. Fig. 4). Surprisingly, *Bt*LolB-depleted cells grew on all provided carbohydrates with minimal defects, the most common being the failure to achieve the same maximum optical density (OD) as wildtype (Fig. 4, Supp. Fig. 2). Perhaps more intriguing, these cells also grew well on carbohydrates that Δ*lnb* cells struggled on the most, namely amylopectin, mucin O-glycans (MOG), and arabinan (Fig. 4B-D).

### Sus lipoproteins localize to the cell surface even upon *Bt*LolB depletion

We previously demonstrated that deletion of Lnb decreases cell surface localization of SusE and SusG, two lipoproteins involved in starch utilization (e.g. amylopectin) (5, 19). We hypothesized that this localization defect at least partially explains the growth lag displayed by Δ*lnb* cells on amylopectin (Fig. 4B) (19). As *Bt*LolB-depleted cells comparatively grow well on amylopectin, we questioned the surface localization of SusE and SusG in these cells. Thus, we immunostained intact cells of wildtype and the *Bt*LolB-inducible strain grown with and without inducer for both Sus lipoproteins (Fig. 5). We also imaged Δ*susE* and Δ*susG* cells as negative controls, and Δ*lnb* cells for a direct comparison. Interestingly, no significant decrease in signal intensity was observed for either SusE or SusG in *Bt*LolB-depleted cells from the wildtype or *Bt*LolB-induced cells, while deletion of Lnb again resulted in less surface presentation of both lipoproteins. (*right*) Montages of representative bacterial cells from the datasets. Cells are outlined in white.

**Figure 4.**
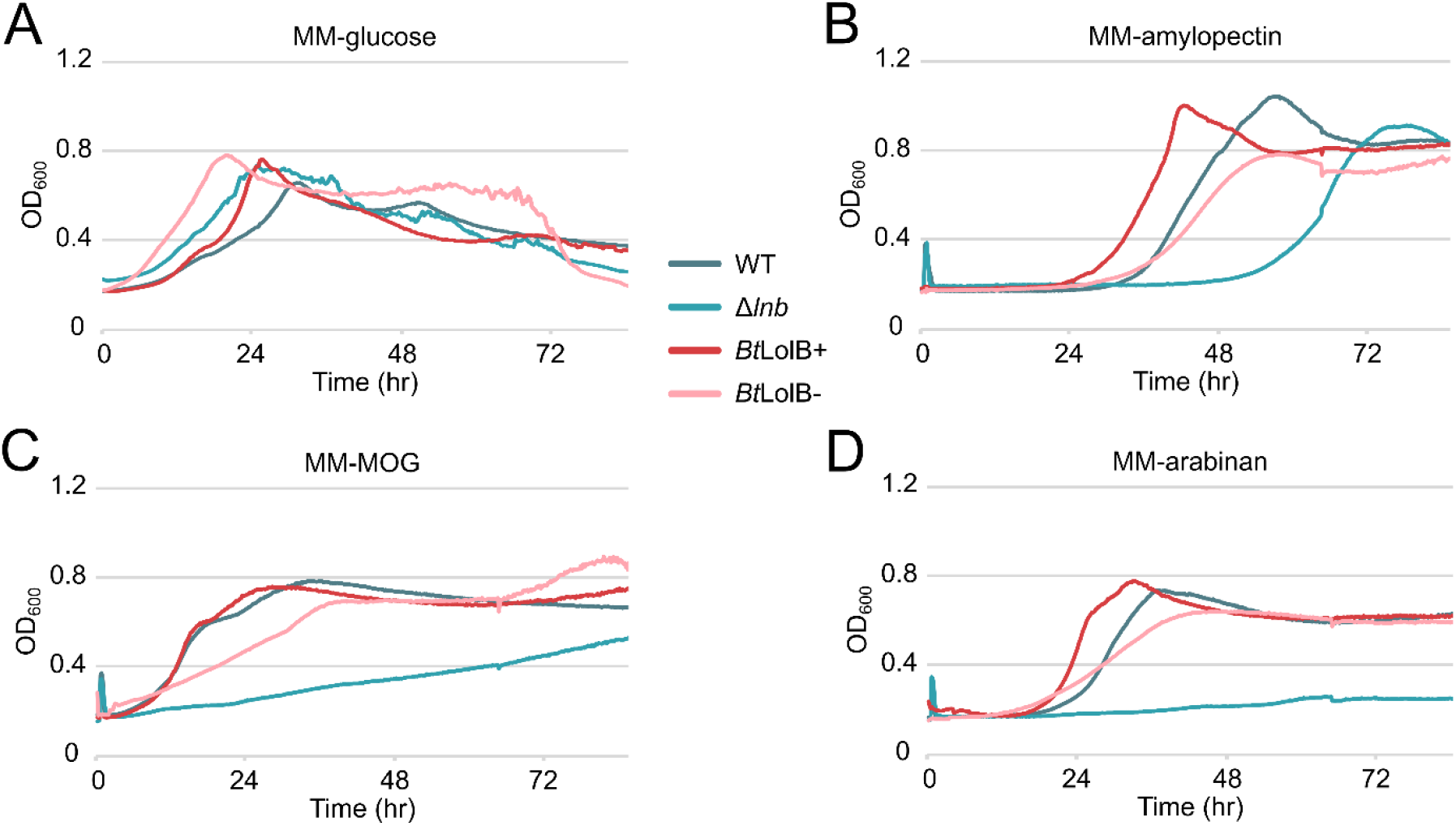
Growth of *B. theta* strains in MM with various carbohydrates. The OD_600_ of *B. theta* wildtype, Δ*lnb*, and the *Bt*LolB-inducible strain grown with and without aTc inducer (respectively indicated as *Bt*LolB+ and *Bt*LolB-) in (A) MM-glucose, (B) MM-amylopectin, (C) MM-MOG, and (D) MM-arabinan over time. Curves are the average of three technical replicates, with an outlier well of Δ*lnb* cells grown in MM-glucose without aTc omitted from the dataset. The graphs shown are representative of three biological replicates.

**Figure 5.**
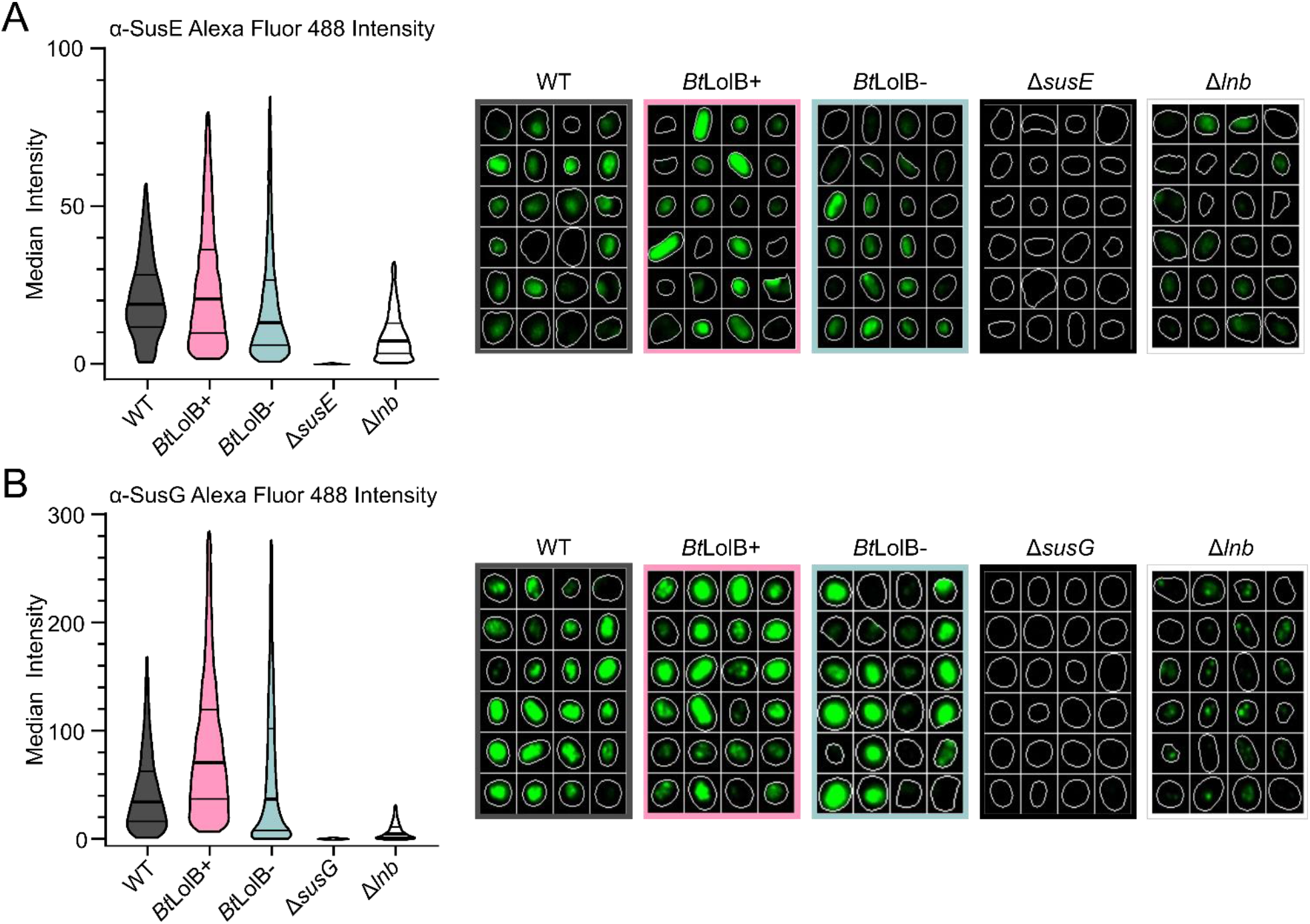
Surface localization of Sus lipoproteins in *B. theta*. Violin plots depict the full distribution of single-cell median fluorescence intensities of the Alexa Fluor 488 signal for the lipoproteins (A) SusE and (B) SusG of the indicated *B. theta* strains. *Bt*LolB+ and *Bt*LolB-refer to the *Bt*LolB-inducible strain KA308 grown with or without aTc, respectively. The width of each violin is proportional to the kernel density estimate of the data (n>1,400 cell observations per condition), and the horizontal lines within each violin indicate the first quartile (25^th^ percentile), median (50^th^ percentile), and third quartile (75^th^ percentile). Montages of representative bacterial cells from the datasets are shown on the right with cells outlined in white.

### Depletion of *Bt*LolB alters the *B. theta* membrane proteome

In *F. johnsoniae*, deletion of LolB results in global changes in protein abundance in the OM fraction (23), and we hypothesized the same would be true for *B. theta*. We grew *B. theta* wildtype and the *Bt*LolB-inducible strain without aTc in TYG, then isolated the IM and OM fractions from each. More NADH consumption was measured in both wildtype and *Bt*LolB-depleted IM fractions than OM fractions, indicating separation of membranes (Supp. Fig. 5). Analysis of the fractions by SDS-PAGE revealed differences in protein profile and abundance of both membranes between wildtype and the *Bt*LolB-depleted samples (Fig. 6A).

**Figure 6.**
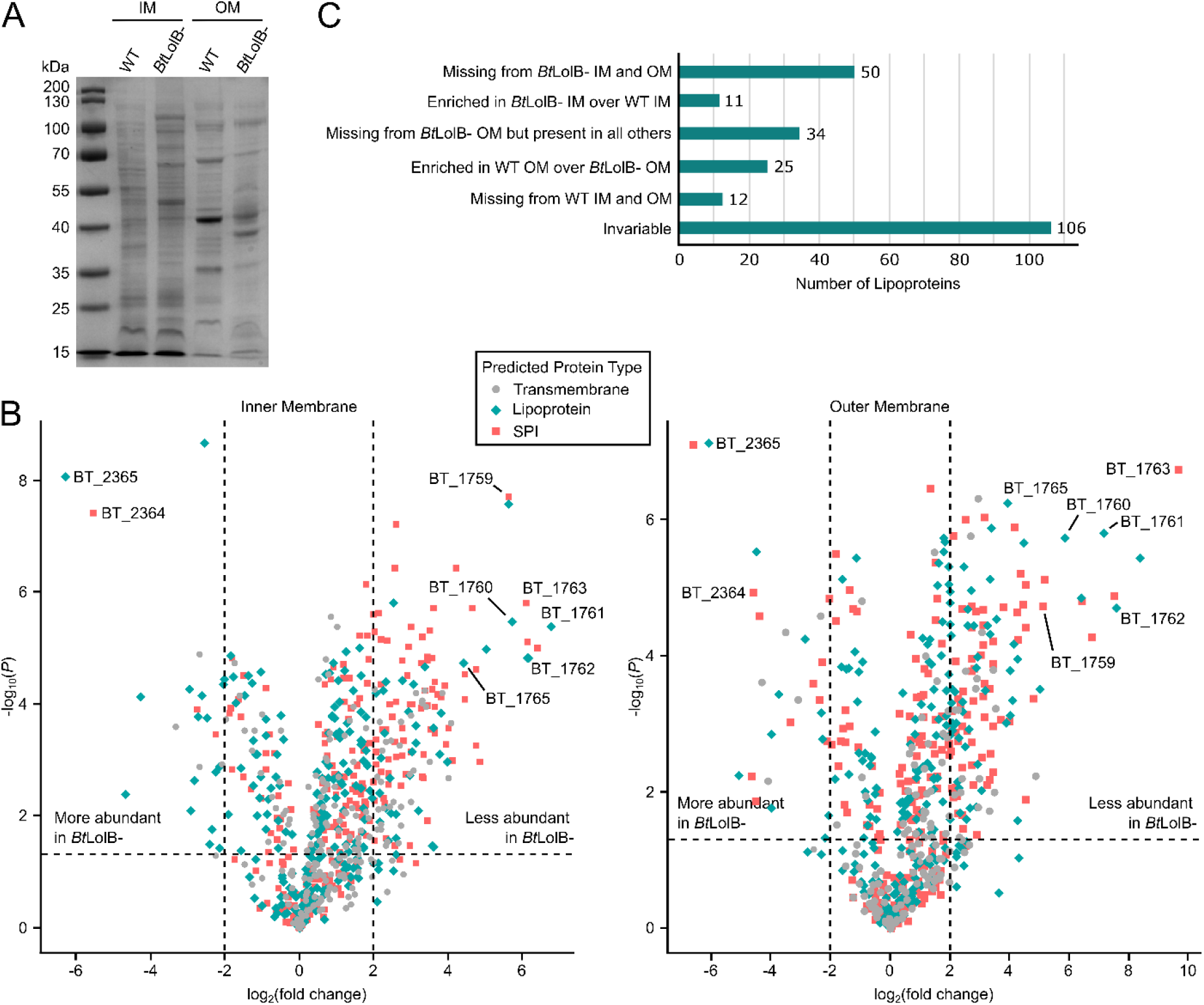
Comparison of membrane fractions of wildtype and *Bt*LolB-depleted *B. theta* cells. (A) Protein profiles of the IM and OM fractions as visualized by SDS-PAGE and Coomassie blue staining. Equal amounts of protein were loaded in each lane (20 µg). (B) Volcano plots showing the inner membrane (*left*) and the outer membrane (*right*) proteomes of wildtype *B. theta* versus *Bt*LolB-depleted samples. Predicted protein types (transmembrane, SPI, or lipoprotein) are indicated by point shape and color, and data points of interest are labelled. The vertical dashed lines represent log_2_(fold change) = ±2.0, and proteins above the horizontal dashed line have *P* < 0.05. (C) A bar graph quantifying lipoproteins into categories of interest. A lipoprotein is “missing” if no intensity was measured for at least 3 of the 4 biological replicates, or it is “enriched” if it is not missing, the log_2_(fold change) is >2.0 or <-2.0, and has a *P* < 0.05.

To define membrane composition, we performed comparative proteomic analysis of the fractions (Supp. Dataset 2). We first noted that no *Bt*LolB was detected in the *Bt*LolB-depleted samples. “Predicted Protein Types” (cytoplasmic, transmembrane, SPI, or lipoprotein (SPII)) were assigned to sort the data, though as we suggested in Table 1, SignalP prediction of SPI versus SPII may not be fully accurate for Bacteroidota. Omitting cytoplasmic proteins, we generated two volcano plots (Fig. 6B). Despite *Bt*LolB’s role in localizing lipoproteins, there was no observable trend in the protein type altered by *Bt*LolB depletion.

Overall, many more proteins are less abundant or missing from the *Bt*LolB-depleted IM and OM fractions than the wildtype fractions. The most altered proteins are BT_1759-BT_1763 and BT_1765, members of an operon involved in fructose and levan utilization that also includes BT_1757 and BT_1758 (42, 47). In our dataset, BT_1759-BT_1762 and BT_1765 are missing from *Bt*LolB-depleted IM and OM fractions, while BT_1763 is missing from the IM and significantly less abundant in the *Bt*LolB-depleted OM compared to wildtype. Another highly altered operon is BT_2364-BT_2365, an uncharacterized SusCD pair, that is more abundant in both *Bt*LolB-depleted membrane fractions over wildtype.

To analyze lipoproteins specifically and represent the data another way, we created a bar graph (Fig. 6C). Across the four fractions (IM and OM of wildtype and *Bt*LolB-depleted cells), 254 total lipoproteins were detected. Fifty are missing from both *Bt*LolB-depleted membrane fractions, potentially suggesting degradation or loss of mislocalized lipoproteins. Eleven lipoproteins are enriched in the *Bt*LolB-depleted IM compared to the wildtype IM, perhaps mirroring the IM accumulation of lipoproteins observed upon LolB depletion in *E. coli* (24). Most strikingly, 34 lipoproteins are missing from the *Bt*LolB-depleted OM but are present in all other fractions, suggesting these lipoproteins specifically could not be inserted into the OM. Another 25 lipoproteins are enriched in the wildtype OM over the *Bt*LolB-depleted OM, suggesting inefficient insertion into the OM. On the other hand, 12 lipoproteins are missing from wildtype fractions but are present in either *Bt*LolB-depleted IM or OM, perhaps gaining importance when *Bt*LolB is lost. Finally, 106 lipoproteins exhibited no significant differences between the samples.

## Discussion

While the Lol system is widely conserved across diderm bacteria, LolB is the least conserved component (22, 34). Recently, structural prediction analyses identified LolB-like proteins throughout Bacteroidota, with some species encoding multiple per genome (23). Here we independently identified the *Bacteroides* LolB via a rescue assay in *E. coli* and validated its lipoprotein insertase activity *in vitro* (Fig. 1 and 3). The mechanism of lipoprotein insertion into the OM by LolB is currently unknown but seemingly involves its protruding loop (32). This loop is conserved in *Bt*LolB and *F. johnsoniae* LolB2, and mutations to the loop abrogate complementation of *Ec*LolB depletion (in the case of *Bt*LolB; Fig. 2B) or hinder gliding motility (for *Fj*LolB2; (23)). Despite their structural similarities, *Fj*LolB2 could not complement the deletion of *Ec*LolB (23), whereas *Bacteroides* LolB complemented *Ec*LolB depletion in our assay.

Depletion of *E. coli* LolB ultimately results in cell death due to toxic accumulation of lipoprotein localization intermediates (e.g. LolA:lipoprotein) in the periplasm and mislocalized lipoproteins in the IM (24). Interestingly, De Smet *et al*. was able to delete the two LolB homologs of *F. johnsoniae*, but not the LolB of *Capnocytophaga canimorsus* (23). We could not delete *Bt*LolB from *B. theta*, but could deplete it to extreme levels, illustrated by the absence of *Bt*LolB in our proteomics dataset (Supp. Dataset 2). It remains unclear why we can deplete but not delete *Bt*LolB, or what determines the genera-level differences in essentiality of LolB within Bacteroidota.

Despite *Bt*LolB depletion, we observed minimal growth defects of *B. theta* on the carbohydrates provided (Fig. 4, Supp. Fig. 2), and no significant decrease in cell surface presentation of Sus lipoproteins (Fig. 5). As capture of carbohydrates by surface lipoproteins is the first step in nutrient uptake by these bacteria, it is possible that *Bt*LolB’s overall impact on growth is negligible. Indeed, our proteomic dataset demonstrates a shuffling in abundance of several SusCD pairs and/or operons upon *Bt*LolB depletion that may compensate for growth in undetermined ways. *Bt*LolB depletion likely results in other phenotypes not investigated in this study, such as changes in lipid composition, capsule and OM vesicle formation, or others.

Nevertheless, our proteomics revealed significant changes to lipoprotein localization in *Bt*LolB-depleted cells (Fig. 6, Supp. Dataset 2). Most interesting are the 34 lipoproteins missing from the *Bt*LolB-depleted OM but are present in all other fractions, suggesting they are canonically *Bt*LolB dependent for OM insertion/localization. The 25 lipoproteins enriched in WT OM over *Bt*LolB-depleted OM, however, indicate that these lipoproteins still reach the OM without *Bt*LolB, albeit to a lesser amount, supporting evidence for an alternate pathway(s) for lipoprotein transport. There are several other known methods for lipoprotein transport, including the Type 2 and Type 5 Secretion Systems, and the surface lipoprotein assembly modulator, but these are missing from Bacteroidota (22, 48, 49). Some bacteria co-opt the lipopolysaccharide transport (Lpt) system for lipoproteins, but it is currently unclear if Bacteroidota do the same (50, 51). Our proteomics did not reveal obvious candidates for an alternate LolB-like protein or lipoprotein transport system, based on gene annotations and AlphaFoldDB structure predictions. It is possible that the protein or system abundance is not significantly altered and thus was not considered. More likely, the protein or system is novel and will not be found by protein “gazing”.

Our previous work identified Lnb, the lipoprotein *N*-acyltransferase in *Bacteroides*, responsible for lipoprotein *N*-acylation (19). Compared to *Bt*LolB depletion, deletion of Lnb results in larger growth deficiencies and less surface presentation of Sus lipoproteins (Fig. 4 and 5, Supp. Fig. 2), suggesting that lipoprotein *N*-acylation is more important than insertion into the OM. This makes sense considering the likelihood of an alternate system for lipoprotein transport to the OM and cell surface, plus the importance of cell surface lipoproteins to *Bacteroides* physiology. Non-*N*-acylated (i.e. diacylated) lipoproteins are considered poor substrates for LolCDE, the complex that extracts lipoproteins from the IM for transport to the OM (20, 52, 53). Luo *et al*. recently demonstrated crosstalk of the Lol and Lpt system in *Pseudomonas aeruginosa*, where lipoproteins are extracted from the IM by LolCDE then transferred to the Lpt system for cell surface localization (51). Should the same be true in *Bacteroides*, then *N*-acylation is a key step for efficient entry into both transport systems. Further studies are required to fully understand lipoprotein transport and how it impacts *Bacteroides* physiology.

## Methods

### Bacterial strains and growth conditions

Strains used in this study are listed in Table S1. *E. coli* strains were grown aerobically in Luria Bertani (LB) medium at 37°C with agitation. When appropriate, cultures were supplemented with 0.2% (wt/vol) L-arabinose or D-glucose. Antibiotic markers were selected with carbenicillin (100 µg/mL), kanamycin (25 µg/mL), and spectinomycin (50 µg/mL).

*Bacteroides* spp. were cultured in a Coy anaerobic chamber with an atmosphere of 10% H_2_, 5% CO_2_, and 85% N_2_ at 37°C. Strains were routinely grown in TYG rich medium (54). Minimal medium was prepared as previously (19), and carbohydrates were added to 5 mg/mL, except MOG (10 mg/mL). Antibiotics were supplemented to 200 µg/mL for gentamicin, 20 µg/mL each for erythromycin and chloramphenicol, and 3 µg/mL for tetracycline. Rhamnose was supplemented to 2.5 mg/mL, 5-fluorodeoxyuridine (FUdR) to 200 μg/mL, and anhydrotetracycline (aTc) to 40 ng/mL. For depletion of *Bt*LolB in strain KA308, no aTc was added to overnight cultures started from glycerol stocks nor any subsequent outgrowths.

### Plasmid construction

Primers and plasmids used in this study are listed in Tables S2 and S3, respectively. The *B. fragilis* genomic library was constructed as previously described (19). The allelic exchange vector pDUDE (<u>D</u>eletion <u>U</u>nder <u>D</u>irected <u>E</u>xchange) was constructed by replacing the tetracycline selection marker *tetQ* from the parent vector pLGB30 (55) with the chloramphenicol resistance marker *catP*. Briefly, *tetQ* was excised using NdeI and SalI (NEB), the latter residing within the multiple cloning site (MCS). The constitutive promoter P1E6 and ribosomal binding site 8 (RBS8) (56) were fused to the *catP* gene to create a construct containing the 5’ NdeI and 3’ SalI sites. Additionally, an SpeI site was included next to the SalI site to increase available restriction sites within the MCS. This construct was ligated into the remaining pLGB30 backbone still harboring the rhamnose-inducible ssBfe1 counterselection system, then whole-plasmid sequenced to verify proper assembly of the vector. Other plasmids were constructed by FastCloning or with the In-Fusion HD Cloning Kit (Takara) (57).

### Construction of the *E. coli P*_*BAD*_*-lolB* strain and *Bacteroides* strains

To create the conditional lethal *lolB* strain of *E. coli*, the arabinose-inducible cassette encoding the kanamycin resistance gene, *rrnB* transcriptional terminator, *araC* gene, and *P*_*BAD*_ promoter was PCR-amplified from KA349 (28) with primers KA39 and KA29. These primers include ∽45-bp homology regions for integration of the cassette upstream of *EclolB* by Red-mediated combination (58). Integrants were selected on arabinose and kanamycin, then verified by PCR. Integration was performed in the *lpp*^*-*^ strain TXM327 (28) to yield strain KA38.

To create strain KA282, *BtlolB-myc* was overexpressed from pWW1376 (56), a pNBU2-based vector that integrates at a chromosomal *att* site. The native *BtlolB* was then deleted via allelic exchange using a variant of the pExchange-*tdk* (59) plasmid conveying tetracycline resistance. For strain KA308, *BtlolB-myc* was cloned into a pNBU2-based vector modified for aTc-inducible expression (46) and integrated into an *att* site. The native *BtlolB* was again deleted by allelic exchange, instead using pDUDE with aTc included in the media throughout the deletion process. pDUDE abides by chloramphenicol selection and rhamnose induction of ssBfe1 toxicity.

### Colony formation by KA38 transformed with experimental plasmids

KA38 cells were grown in LB plus kanamycin and arabinose to an OD_600_ of ∽0.5 to 0.6, then electroporated with 50 ng of plasmid. After 1 hr outgrowth in just LB, 100 μL each was spread onto three LB agar plates containing arabinose and three containing glucose. All plates contained carbenicillin for transformant selection. Colonies were enumerated the following day. Three biological replicates were performed, each with three technical replicates.

### Growth analysis

For *E. coli*, overnight cultures were washed thrice in LB to remove arabinose. Cells were diluted 1:6,000 into fresh LB containing carbenicillin and arabinose or glucose in a 96-well plate. The OD_600_ was measured every 10 min in a BioTek LogPhase 600 microbiology reader at 37°C with constant shaking at 800 rpm. For *B. theta*, overnight cultures were washed thrice in PBS, then normalized to a starting OD_600_ of 0.05 in the intended media and carbon source in a 96-well plate. The OD_600_ was measured every 10 minutes in a BioTek automated plate reader at 37°C. Curves shown are the average of three technical replicates and are representative of three biological replicates.

### Purification of LolA:Lpp complexes

Overnight cultures of *E. coli* strains KA315 and KA316 constitutively expressing Lpp(K58A)-Strep were inoculated 1:100 into fresh LB containing spectinomycin and kanamycin. Cells grew at 37°C until reaching an OD_600_ of ∽0.2, then expression of *Ec*LolA-His (KA315) or *Bt*LolA-His (KA316) was induced by adding IPTG to 0.5 mM. Cultures were harvested at an OD_600_ of ∽1.0. To release periplasmic material, the pellets were resuspended in 1 mL of 100 mM Tris-HCl (pH 8.0), 500 mM sucrose, 1 mM EDTA, and 1 mg/mL lysozyme, then incubated on ice for 30 min. Tubes were spun at 21,100 x g for 30 min at 4°C to pellet cell debris. The resulting supernatant was mixed with ∽250 µL of HisPur Ni-NTA Resin (Thermo Scientific) equilibrated with Buffer A (25 mM NaH_2_PO_4_, 500 mM NaCl, 20 mM imidazole, pH 7.4) and flipped for 1 hr at 4°C. The bound resin was washed thrice with 1 mL of Buffer A, then once with 1 mL of Buffer A with 50 mM imidazole. The LolA:Lpp complex was eluted with 500 µL of Buffer B (Buffer A with 300 mM imidazole).

### Membrane fractionation

Cells were grown overnight in TYG. Pellets were resuspended in 50 mM HEPES pH 7.4 supplemented with cOmplete EDTA-free protease inhibitor tablet (Roche), then lysed by four passes through an EmulsiFlex-C3 homogenizer (Avestin, Inc.) at 20,000-25,0000 psi. Unbroken cells were removed by centrifugation at 5,000 x g for 5 min at 4°C. Total membranes were collected from the supernatant by ultracentrifugation in a swinging bucket rotor at 30k rpm for 1.5 hr at 4°C. For the lipoprotein insertion assay and co-immunoprecipitation, the total membranes were solubilized overnight by flipping at 4°C in 12 mL of HEPES buffer with 1% (wt/vol) sarcosine and protease tablet. Samples were ultracentrifuged again to collect the OM fraction, which was then solubilized overnight by flipping at 4°C in 1 mL of HEPES buffer with 1% (wt/vol) n-dodecyl β-D-maltoside (DDM). For collection of IM and OM fractions for proteomic analysis, total membranes were instead solubilized in 1 mL of the HEPES buffer with sarcosine. Following ultracentrifugation at 100,000 x g for 1 hr at 4°C, the supernatant was saved as the IM fraction, and the OM pellet was resuspended in 1 mL of HEPES buffer. Protein concentrations were determined by BCA Protein Assay Kit (Thermo Scientific), then lyophilized.

### Lipoprotein insertion assay

0.75 mg/mL of OM fraction was combined with 20 µL of the purified *Ec*LolA-His:Lpp-Strep or *Bt*LolA-His:Lpp-Strep in a 100 µL reaction volume (brought up with LB). The reactions were incubated for 1 hr at 37°C, then centrifuged at 21,100 x g for 20 min to pellet the OM. Pellets were rinsed with PBS to remove residual LolA:Lpp, then resuspended in 30 µL PBS. Ten µL of the resuspension was loaded onto a 15% Tris-glycine gel to detect Lpp-Strep and onto 10% Tris-glycine gels to detect LolA-His.

### Immunoblotting

Samples were separated by SDS-PAGE, then transferred to 0.2 µm polyvinylidene difluoride (PVDF) membrane. Membranes were incubated with a 1:5,000 dilution of StrepTactin-horseradish peroxidase (HRP) conjugate (Bio-Rad) or a 1:500 dilution of 6x-His Tag Monoclonal Antibody HRP conjugate (Invitrogen). Signals were detected by enhanced chemiluminescence.

### RNA isolation and qPCR

Overnight cultures were OD_600_-normalized to 0.05, then grown to an OD_600_ of 1.0 in 5mL of TYG. An equal volume of RNAprotect (Qiagen) was added, incubated for 5 min, then centrifuged at 8.8 k x g for 10 min. The supernatant was decanted and pellets stored at -80°C until extraction. For RNA isolation, cells were resuspended in TE buffer containing 1 mg/mL lysozyme, then mixed with RLT buffer supplemented with β-mercaptoethanol according to the RNeasy Kit protocol (Qiagen). RNA was extracted twice with phenol:chloroform:isoamyl alcohol, followed by an extraction with just chloroform. RNA was purified via standard sodium acetate/isopropanol precipitation, then treated with TURBO DNA-*free* (Invitrogen) following manufacturer’s instructions. RNA was re-purified by precipitation, resuspended in RNase-free water, and quantified by NanoDrop (Thermo Scientific). For qPCR, reverse transcription was performed using SuperScript III reverse transcriptase and random primers (Invitrogen). The resulting cDNA was quantified using a homemade qPCR mix as previously described (60) and ran for 40 cycles of 95°C for 3 sec, 55°C for 20 sec, and 72°C for 20 sec in a CFX Connect Real-Time thermocycler (Bio-Rad). A melt curve was generated to measure amplicon purity. Using the ddCT (delta-delta Cycle Threshold) method, the data was 16S rRNA-normalized and compared to the wildtype strain to calculate the fold change. qPCR was performed in duplicate with RNA isolated from three biological replicates.

### Formaldehyde cross-linking

The *Bt*LolB-Myc overexpression strain KA282 was crosslinked with formaldehyde as described previously (61). Briefly, cells pelleted from overnight cultures were washed twice with 0.1 M potassium phosphate pH 7.2, then resuspended in ½ volume of buffer supplemented with cOmplete EDTA-free protease inhibitor tablet (Roche). Formaldehyde was added to 1% and samples were incubated for 1 hr at room temperature. Cells were washed twice with potassium buffer, then pelleted and stored at -80°C until use.

### Co-immunoprecipitation

*Bt*LolB-Myc was immunoprecipitated using Pierce Magnetic Beads (Thermo Scientific) for Anti-c-Myc or Anti-HA and a MagJET Separation Rack (Thermo Scientific). One mL of DDM-solubilized OMs per replicate was divided in half, then mixed with 25 µL Anti-HA or Anti-c-Myc beads that had been washed twice with 1 mL PBS. The mixtures were flipped for 1 hr at room temperature. Bound beads were washed thrice with PBS followed by acetone precipitation. Four replicates were performed.

### NADH oxidase assay

Based on previous protocols (62, 63), 60 µg/mL of membrane fraction was mixed with buffer containing 50 mM Tris-HCl pH, 0.2 mM dithiothreitol (DTT), and 0.12 mM NADH in a 2 mL volume. Reactions were incubated anaerobically at 37°C. Absorbance at 340 nm was measured at 0, 24, and 48 hr.

### Proteomic analysis

Lyophilized samples were solubilized in 4% SDS, 100 mM HEPES by boiling for 10 min at 95°C, then the protein concentrations of cell fractions measured using bicinchoninic acid protein assays (Thermo Fisher Scientific) with 100 µg of each fraction utilized for analysis. Immunoprecipitated protein samples were solubilized in 4% SDS, 100 mM HEPES by boiling for 10 min at 95°C and assumed to contain <50 µg of protein. Both membrane fractions and immunoprecipitated samples were cleaned-up using S-trap mini columns (Protifi, USA) according to manufacturer’s instructions. Samples were reduced with 10 mM DTT for 10 min at 95°C, then alkylated with 40 mM iodoacetamide for 1 hr in the dark. Samples were acidified to 1.2% with phosphoric acid and diluted with seven volumes of S-trap wash buffer (90% methanol, 100 mM tetraethylammonium bromide pH 7.1) before being loaded onto S-traps then washed thrice with 400 µL S-trap wash buffer. Samples were digested with a 1:50 protease:protein ratio of trypsin (4 µg for fractions and 1 µg for immunoprecipitated samples) in 100 mM tetraethylammonium bromide overnight at 37°C before collection by centrifugation, followed by sequential washes with 100 mM tetraethylammonium bromide, 0.2% formic acid, then 0.2% formic acid/50% acetonitrile. Samples were dried down and further cleaned up using C18 Stage (64, 65) tips to ensure removal of particulate matter.

### Reverse phase liquid chromatography-mass spectrometry

C18-purified peptide samples were resuspended in Buffer A* (2% acetonitrile, 0.1% trifluoroacetic acid in Milli-Q water) and separated using a two-column chromatography setup on a Dionex Ultimate 3000 UPLC composed of a PepMap100 C18 20 mm × 75 µm trap and a PepMap C18 500 mm × 75 µm analytical column (Thermo Fisher Scientific) coupled to an Orbitrap Fusion™ Lumos™ Tribrid™ Mass Spectrometer (Thermo Fisher Scientific) with a FAIMS Pro interface (Thermo Fisher Scientific). 95-minute gradients were run for each sample, with samples loaded onto the trap column with 98% Buffer A (2% acetonitrile, 0.1% formic acid in Milli-Q water) and 2% Buffer B (80% acetonitrile, 0.1% formic acid) with peptides separated by altering the buffer composition from 2% Buffer B to 28% B over 76 min, from 28% B to 40% B over 9 min, then from 40% B to 80% over 3 min. The composition was held at 80% B for 2 min, dropped to 2% B over 2 min, then held at 2% B for another 3 min. The Orbitrap Lumos Mass Spectrometer was operated in a data-dependent mode, switching between the collection of a single Orbitrap MS scan (300-1600 m/z at a resolution of 60k) every three seconds followed by the collection of Orbitrap MS/MS HCD scans of precursors (fixed NCE 35%, maximal injection time of 54 ms, and a resolution of 15k).

### Proteomic data analysis

Datafiles were searched against the *Bacteroides thetaiotaomicron* VPI-5482 proteome (UniProt: UP000001414) using MaxQuant (V2.2.0.0)(66) allowing carbamidomethylation of cysteine set as a fixed modification and oxidation of methionine as a variable modification. Searches were performed with trypsin cleavage specificity, allowing two miscleavage events with a maximum false discovery rate (FDR) of 1.0% set for protein and peptide identifications. The LFQ and “Match Between Run” options were enabled to allow comparison between samples. The resulting data files were processed using Perseus (v1.4.0.6)(67) with missing values imputed based on the total observed protein intensities with a range of 0.3 σ and a downshift of 1.8 σ. Statistical analysis was undertaken in Perseus using two-tailed unpaired t-tests. Predicted protein types (cytoplasmic, transmembrane, SPI, or lipoprotein (SPII)) were assigned using TOPCONS and SignalP as done by others (19, 26, 38, 68).

### Immunostaining, image acquisition, and single-cell analysis

Formalin-fixed cells were prepared for imaging with anti-SusG or -SusE antibodies and Alexa Fluor 488 secondary antibody as previously described (19). Cells were incubated for 30 min in the dark with Hoechst-33342 stain (Thermo Fisher Scientific) diluted to 10µg/mL with PBS. Wells of a flat, clear-bottomed 96-well plate (Revvity PhenoPlate Cat. No. 6055600) were coated with 0.01% poly-ʟ-lysine and washed 2-3 times with PBS. Cells were deposited to achieve sufficient cell density for imaging (70 or 100 µL of cells, either undiluted or diluted 1:10 in PBS) per well and were incubated at room temperature or 37°C for 30-60 min in the dark. Images were acquired and analyzed as previously described (19) using a Yokogawa CQ1 high-content imaging instrument. Cells were imaged with a 60x/0.9 NA dry objective lens and Hoechst images were taken in the blue channel with an excitation wavelength of 405 nm with 1 sec exposure times. Between 80-180 fields per well were imaged to generate approximately 10,000 cell observations. Segmentation and feature extraction was performed with CellProfiler 4.2.8 using the Hoechst channel for segmentation and intensity and size/shape features were computed for the Alexa Fluor 488 and Hoechst channels. Three independent biological replicates were performed for SusG and one for SusE.

## Data availability

The mass spectrometry proteomics data has been deposited in the Proteome Xchange Consortium via the PRIDE partner repository with the data set identifier: Project accession: PXD073329 (Username: Password: BePmBthjFECJ) and Project accession: PXD073339 (Username: Password: LrdfOQtmstDA).

## Author contributions

K.M.A.: conceptualization, investigation, project administration, visualization, writing – original draft. R.T.: investigation, methodology, visualization, writing. N.E.S.: investigation, writing. N.A.P.: methodology, supervision. J.W.W.: formal analysis. J.Z.S.: visualization, writing, resources. N.M.K.: conceptualization, funding acquisition, project administration, supervision, visualization, writing.

## Acknowledgements

This work was supported by a University of Michigan Elizabeth Crosby Award and NIH grant [1R21AI180287](https://public.era.nih.gov/grantfolder/viewCommonsStatus.era?applId=10784857) to N.M.K. J.Z.S. was supported by NIH S10 OD034245 and R01 GM152417. N.E.S. was supported by an Australian Research Council Future Fellowship (FT200100270), an ARC Discovery Project Grant (DP210100362) and a NHMRC Ideas Grant (GNT2018980). We thank the Melbourne Mass Spectrometry and Proteomics Facility of The Bio21 Molecular Science and Biotechnology Institute for access to MS instrumentation.

